# Aortic valve stenosis promotes pathological shear stress-dependent epigenomic dysregulation in circulating T cells

**DOI:** 10.64898/2026.05.17.722103

**Authors:** Yunyang Zhang, Jan Lucas Kleiner, Jimmy Zheng, Frank Splettstoesser, Sebastian Zimmer, Mark Coburn, Christina Weisheit, Stilla Frede, Mark E. Pepin

## Abstract

**Background:** Calcific aortic valve stenosis (AVS) is the most prevalent valvular heart disease in Western adults, yet no disease-modifying therapy exists. High shear stress (HSS) generated by progressive valvular obstruction drives endothelial injury and immune-mediated inflammation, but the contribution of circulating T cells to AVS pathogenesis remains poorly defined.

**Objectives:** We tested whether chronic HSS corresponds with epigenomic reprogramming of peripheral T cells proportionate with hemodynamic severity to yield a clinically informative proxy of disease.

**Methods:** A prospective cohort of 70 participants was recruited for peripheral blood sampling, including 34 with severe symptomatic AVS (aortic valve area <1.0 cm^2^, mean gradient ≥40 mmHg) scheduled for transcatheter aortic valve implantation and 36 age- and sex-matched controls. Peripheral T cells were isolated and profiled by genome-wide CpG methylation (Illumina MethylationEPIC) and RNA-sequencing. To test whether HSS directly activates inflammatory signaling, Jurkat T cells were exposed to 20 dyn/cm^2^ HSS via parallel-plate microfluidic chamber and concomitant CD3/CD28 stimulation, followed by assessment of NFAT nuclear translocation and NFAT target gene expression.

**Results:** Unsupervised clustering of the 5,000 most-variable CpG loci resolved an epigenomic axis segregating AVS from control T cells (PC1, 15.8% variance explained; *P* = 3.9×10^-6^). Multivariable-adjusted analysis identified 3,950 differentially methylated positions (1,889 hyper-, 2,061 hypo-methylated), enriched in promoter-associated CpG islands implicating aortic valve morphogenesis (P = 6.0 x 10^-10^) and cell-cell adhesion pathways (P = 9.5 x 10^-5^). Multi-omics factor analysis isolated a latent factor that independently associated with AVS (adjusted P = 1.8×10^-3^; AUC = 0.79), enriched for chemokine receptor binding and TNF-family signaling, and correlated with canonical HSS-responsive transcripts, consistent with a T cell-mediated shear stress activation. An 18-CpG elastic-net methylation risk score discriminated AVS from controls (AUC = 0.89) and independently predicted hemodynamic severity (β = 7.05 mmHg/SD, 95% CI 2.31-11.79). HSS augmented NFAT nuclear translocation in CD3/CD28-activated Jurkat T cells and induced NFAT-responsive inflammatory transcripts.

**Conclusions:** Severe AVS is associated with promoter-enriched epigenomic remodeling of circulating T cells that converges on hemodynamic stress-dependent inflammatory programs. An 18-CpG methylation risk score outperforms clinical covariates and tracks hemodynamic severity, establishing peripheral T cell DNA methylation as a molecular corollary of AVS.

## Introduction

Calcific aortic valve stenosis (AVS) is the most prevalent valvular heart disease in elderly adults in Western societies and its incidence is rising sharply with global population aging.^1,2^ Despite decades of investigation, no pharmacologic therapy has yet shown efficacy in forestalling disease progression, and structural aortic valve replacement remains the only definitive treatment for severe or symptomatic disease.^1,3^ Developing novel therapeutic approaches for AVS requires better mechanistic resolution of the biological drivers underlying its pathogenesis.

AVS is increasingly recognized as a chronic inflammatory disorder rather than a passive degenerative process.^4^ Progressive leaflet thickening and calcification generates pathologically elevated transvalvular pressure gradients that subject the valvular endothelium to rising hemodynamic shear stress (HSS).^3,5^ HSS disrupts endothelial integrity, promotes subendothelial lipid infiltration, and upregulates adhesion molecules that facilitate immune cell recruitment, while simultaneously driving valvular interstitial cell (VIC) myofibroblast and osteoblast-like transcriptional trajectories.^4,5^ The resulting cascade of endothelial injury, immune activation, and calcification comprises a pathological succession that remains poorly understood on a gene-regulatory level.

T lymphocytes are emerging as key cellular mediators of HSS-dependent inflammatory processes. Histological analyses of stenotic valvular tissue demonstrate dense T cell infiltration in calcific regions, with both CD4^+^ and CD8^+^ subsets exhibiting activated, tissue-resident memory phenotypes and clonal T cell receptor expansions consistent with antigen-driven responses.^6,7^ Functionally, CD8^+^ T cells impair osteoclast-mediated calcium resorption through IFN-γ secretion, while CD4^+^ T cells promote VIC osteoblastic differentiation via TNF-α and TGF-β, thereby directly accelerating valvular calcification.^8^ Emerging single-cell T cell receptor sequencing data further implicate shared clonal expansions between valvular and circulating T cell compartments,^9^ suggesting that peripheral T cells may serve as peripheral surrogates of the immune state within the diseased valve.

While immune activation in AVS is well-documented, the mechanism by which pathological hemodynamic forces translate into sustained T cell pro-inflammatory programming has only recently been identified. A mechanosensitive ion channel, Piezo1, is constitutively expressed on T cells and functions as a primary transducer of extracellular mechanical stimuli including fluid shear stress.^10,11^ Piezo1 activation by HSS drives rapid Ca^2+^ influx, calcineurin-mediated dephosphorylation, and nuclear translocation of the nuclear factor of activated T cells (NFAT), a master transcriptional regulator of T cell activation, cytokine production (IL-2, TNF-α, IFN-γ), and clonal expansion via Cyclin D1-dependent cell cycle progression.^10-13^ In the setting of severe AVS, circulating T cells are repeatedly exposed to supraphysiologic HSS, thus providing a sensible mechanism by which pathological hemo-dynamics may propagate chronic T cell immune activation beyond endothelial adhesion alone.

Epigenetic modifications, particularly methylation at CpG dinucleotides, are now understood to encode durable transcriptional responses to physiological and environmental stimuli in immune cells, including T cell activation state, effector differentiation, and inflammatory commitment.^14,15^ Growing evidence from epigenome-wide association studies implicates specific CpG methylation alterations in cardiovascular disease, atherosclerosis, and valvular pathology.^16^ However, neither the pathophysiological relevance nor the clinical importance of these epigenomic alterations has been shown in AVS.

Therefore, in the current study we isolated circulating T cells from a prospective cohort of AVS to determine whether reprogramming of CpG methylation (1) accompanies the elevated HSS seen in severe AVS and (2) is associated with AVS disease severity. Using a combination of clinical and *in vitro* experimental designs, we demonstrate that a distinct immuno-epigenomic HSS signature exists that can be distilled into a clinically discriminative methylation risk score (MRS) for AVS. These findings support circulating T cell epigenomics as a novel mechanistic and translational avenue for valvular disease.

## Methods

### Ethics statement regarding use of human tissue

This study was approved by the Ethics Committee of the University Hospital Bonn (Project #AZ078/17), and all participants provided written informed consent in accordance with the Declaration of Helsinki.

### Data availability and open-sourced bioinformatic analysis

Raw and processed files for RNA-sequencing and DNA methylation analyses are made available upon reasonable request. A detailed description of the bioinformatic workflow, including all coding scripts and quality control metrics, are provided as an open-source *GitHub* repository: https://github.com/mepepin/AVS_Epigenomics. We created a reproducible analytic pipeline for this study to define the functional role of CpG methylation both as a proxy of gene regulatory alterations and as a clinical tool to define pathological shear stress-dependent reprogramming of circulating T cells in aortic valvular stenosis (**Figure 1**).

**Figure 1.**
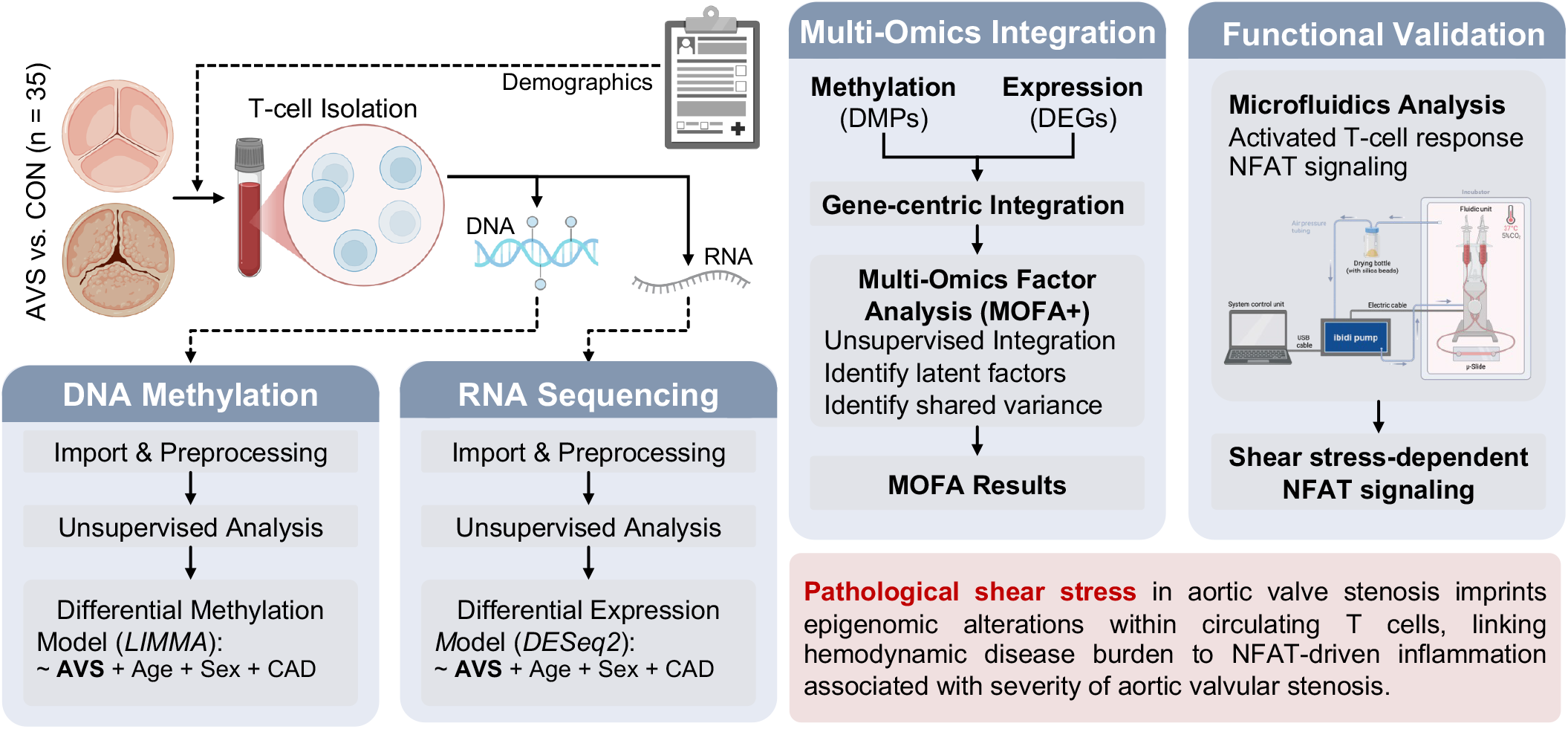
Conceptual overview and computational workflow. Study design overview depicting the isolation of peripheral T cells from 70 participants, including 34 patients undergoing TAVI for severe aortic valve stenosis. DNA and RNA were co-isolated for parallel multi-omics analysis including genome-wide CpG methylation profiling and RNA-sequencing. Integrative analysis via MOFA and elastic-net regression identified functional epigenomic programs and a methylation severity score.

### Tissue biopsy selection and data procurement

We prospectively enrolled a total of 70 patients presenting to the University Hospital Bonn from April 2022 to March 2023. The study cohort consisted of 34 patients with severe, symptomatic aortic valve stenosis (AVS) scheduled for transcatheter aortic valve implantation and 36 age- and sex-matched controls (CON) with preserved aortic valve hemodynamics. Severe AVS was defined according to current ESC/EACTS guidelines as an aortic valve area (AVA) < 1.0 cm^2^ and a mean transvalvular pressure gradient ≥40 mmHg. CON comprised of healthy volunteers with no history of cardiovascular disease.

To minimize confounding effects of immunologic comorbidity, subjects were excluded if they had (1) active systemic infection or sepsis (C-reactive protein > 10 mg/L), (2) known autoimmune or chronic inflammatory disorders, (3) active malignancy, or (4) a history of organ transplantation.

### Sample acquisition and T cell isolation

Peripheral blood mononuclear cells (PBMCs) from patients undergoing transaortic valvular intervention and healthy donors were isolated from whole EDTA blood using Histopaque 1077 and 1119 (Sigma-Aldrich, Germany). Following gradient centrifugation T cells were isolated by using Pan T Cell Isolation Kit (Miltenyi, Germany) according to manufacturer’s instructions. The flow-through was collected as the purified T cell isolate.

### Shear stress assay

10^6^ cells/mL Jurkat cells were obtained from ATCC (American Type Culture Collection, Manassas, VA, USA) and cultured in RPMI 1640 media with 10% FCS in an incubator at 37°C with 5% CO2. Cells were treated with or without antibodies targeting CD3 (OKT3) and CD28 (CD28.2) at a concentration of 2 μg/mL for both antibodies (ThermoFisher, Germany). To determine the impact of HSS on isolated T cells, we subjected Jurkat cells to parallel-plate flow chamber associated shear stress using the Ibidi microfluiding system (µ-Slide VI 0.1 and VI 0.4, ibidi, Germany). The constant flow rates were achieved using a computer-controlled air pump to produce constant wall shear stress (*τ*) defined as follows:

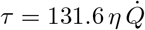

Where τ = wall shear stress (Pa), η = viscosity (Pa·s), and 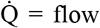 rate (mL/min); the empiric constant (131.6) was provided by the manufacturer based on the microfluidics chamber design. Cell viability was assessed before and after the treatment by acridine orange/ propidium iodide dye. At 3 hours of HSS, Jurkat T cells showed no significant decrements in cell viability (**Supplemental Figure S1**). Subcellular fractionation and Jess capillary immunoassay for NFAT were performed after 1 hour of shear stress to capture early nuclear translocation, whereas downstream NFAT target gene expression (qPCR) was assessed at 3 hours.

### Subcellular Fractionation

Intranuclear protein extracts were prepared using the NE-PER™ Nuclear and Cytoplasmic Extraction Reagents (Thermo Fisher, Germany) per manufacturer’s instructions. Protein concentration was determined using the bicinchoninic acid (BCA) assay (Pierce™ BCA Protein Assay Kit, Thermo, Germany) and bovine serum albumin as a standard according to the manufacturer’s instructions.

### Protein level quantification of NFATc1

NFATc1 was quantified by automated capillary immunoassay on the Jess system (ProteinSimple) using the 12-230 kDa separation module, following the manufacturer’s protocol as previously described.^17,18^ Jurkat lysates were prepared at 0.3 µg/µl in 0.1X sample buffer with fluorescent master mix and denatured at 95 °C for 5 min. NFATc1 (sc-7294, Santa Cruz) and β-actin (8457s, Cell Signaling) were probed in the first cycle and detected with HRP- and NIR-conjugated secondaries, respectively; Lamin A (ab8980, Abcam) was probed after RePlex stripping and detected with an HRP-conjugated secondary. Signal was quantified from area under the curve using Compass software. β-actin and Lamin A served as fractionation controls, and samples with cross-contamination between cytosolic and nuclear fractions were excluded.

### RNA expression analysis

Total RNA was isolated from patients’ T cells and Jurkat cells using NucleoSpin TriPrep (Machery Nagel, Germany), according to the manufacturer’s instructions. Amount and quality of samples were measured by absorbance spectroscopy (Nanodrop Products, Germany). RNA was reverse transcribed using high-capacity cDNA Reverse Transcription Kit (Thermo Fisher Scientific, Germany). Expression of specific target genes was analyzed using validated TaqMan probes and TaqMan Universal Master Mix (Thermo Fisher Scientific, Germany) on a QuantStudio5 (Thermo Fisher Scientific, Germany). Relative expression levels were obtained after normalization to 18S as the expression of this house keeping gene is not modulated by shear stress using the 2^-ΔΔ^ CT method.

### Genome-wide DNA methylation analysis

DNA was extracted from human samples using the NucleoSpin TriPrep (Machery Nagel, Germany), according to the manufacturer’s instructions. 1200 ng DNA was used for the Infinium Methylation EPIC Bead Chip Array (Illumina, San Diego, USA). The EPIC array used in this cohort interrogates over 850,000 CpG sites distributed across promoters, gene bodies, CpG islands, shores, shelves, enhancers, and other regulatory regions.

Methylation array data were used to quantify the proportion of methylation at each CpG locus on a probe-wise basis using the R package *minfi* (1.36.0). Array intensity data were preprocessed using functional normalization to correct for technical differences, as described.^19^ Total (methylated + unmethylated) signal intensity for each probe was weighted against the background signal via negative control probes to provide a statistical (*P*-value) detection threshold; no outliers were identified using this approach. CpG sites overlapping known SNPs at single-base extension or CpG positions and chromosome X/Y probes were removed, retaining 816,126 autosomal CpG probes across 70 samples. Differential methylation analysis used probe-wise linear models fitted to M values (logit-transformed β) in *limma* with empirical Bayes variance shrinkage.^20^ The design matrix included Group (AVS vs. Control) along with CAD, age, and sex for multivariable adjustment. Differentially methylated positions (DMPs) were nominally defined at *P* < 0.05 and |Δβ| > 5%.

### RNA-Sequencing

RNA was isolated from circulating T cells using the NucleoSpin TriPrep (Machery Nagel, Germany) and validated to ensure RNA quality via fragment analysis (Agilent). Samples achieving RNA Integrity Numbers (RINs) > 7 were retained, resulting in 54 patient-derived samples for downstream RNA-sequencing via paired-end 100bp RNA sequencing at Life and Brain GmbH (Bonn, Germany). Prior to alignment, adapters and low-quality (PHRED < 20, or 1% sequencing error rate) sequences were trimmed from read files using *cutadapt* (v5.2). Paired-end 100 bp sequencing was performed at Life and Brain GmbH (Bonn, Germany). Adapters and low-quality bases (Phred < 20) were trimmed using cutadapt (v5.2). Reads were aligned to hg19 using STAR (v2.7.11b; ∼95% uniquely mapped reads). Raw counts were generated using Samtools, and differential expression was performed using DESeq2 (v1.50.2) in R (v4.5.2) with Wald test statistics and Benjamini-Hochberg (BH) adjustment. At BH-adjusted padj < 0.05, 75 DEGs were identified; at nominal *P* < 0.05, 1,677 genes showed differential expression.^21^

### CpG methylation severity score

Mean aortic valve gradient was modeled from clinical covariates (age, sex, smoking, BMI, beta-blocker, ACE-I/ARB) in AVS patients with complete data; severity residuals were screened against CpG M-values by Spearman correlation. The top 100 CpGs were entered into elastic-net Gaussian regression (α = 0.5, λ selected by 5-fold cross-validation). The AVS cohort (n = 29) was then split 1:1 into 15 training and 14 held-out test samples and stratified by mean-gradient severity tertile. The resulting MRS, which retained 18 non-zero CpG coefficients, was projected into the full 70-sample case-control dataset for discrimination analysis.

### Statistics

Normality was assessed by Shapiro-Wilk test. Parametric distributions were evaluated using student t-test with BH adjustment; otherwise, a Mann-Whitney U test was used. Categorical variables were compared by Fisher’s exact test. All data are reported as mean ± SD unless otherwise specified.

## Results

### Baseline Characteristics and Cohort Demographics

To isolate the epigenetic signal of valvular hemodynamics from demographic covariates, we enrolled a prospective cohort of 70 patients, comprising 34 individuals with severe Aortic Valve Stenosis (AVS) scheduled for TAVI and 36 controls with no history of aortic valvular stenosis (**Table 1**). The AVS cohort was significantly older than the control group (80.6 ± 5.2 vs. 57.5 ± 17.3 years; *P* < 0.001) and comprised a higher proportion of females (56% vs. 28%; *P* = 0.017), prompting adjustment for age and gender in the downstream analysis. As expected, the AVS group exhibited significantly reduced aortic valve area (0.8 ± 0.2 vs. 1.5 ± 0.0 cm^2^; *P* = 0.021) and increased mean trans-aortic gradient. Because coronary artery disease (CAD) is a known causal factor in AVS pathogenesis, downstream methylation and RNA-seq analyses retained CAD as a model covariate.

**Table 1:**
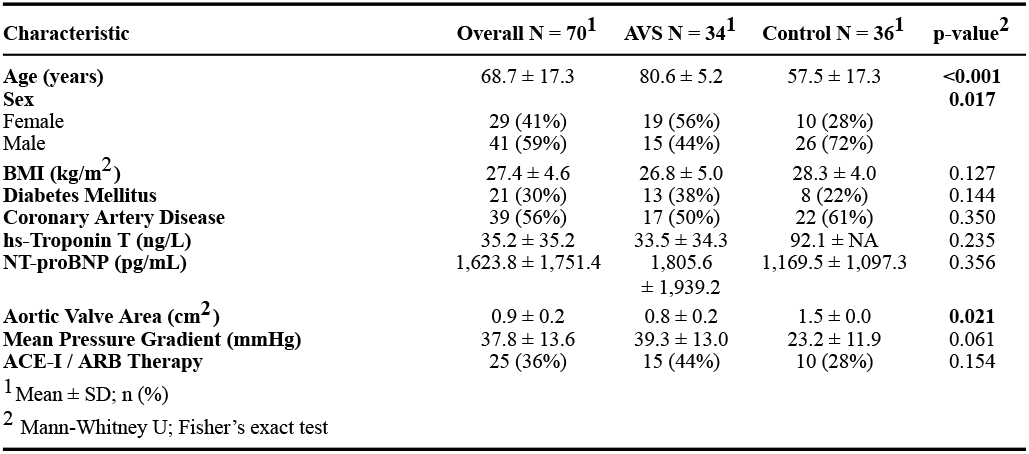
Baseline Clinical Characteristics. Data are mean ± SD or n (%). P-values compare AVS vs. Control groups using Mann-Whitney U (continuous) or Fisher’s exact test (categorical). ACE-I/ARB, angiotensin-converting enzyme inhibitor/receptor blocker; AVA, aortic valve area; BMI, body mass index; CAD, coronary artery disease; NT-proBNP, N-terminal pro-B-type natriuretic peptide.

| Characteristic | Overall N = 70<sup>1</sup> | AVS N = 34<sup>1</sup> | Control N = 36<sup>1</sup> | p-value<sup>2</sup> |
| --- | --- | --- | --- | --- |
| Age (years) | 68.7 ± 17.3 | 80.6 ± 5.2 | 57.5 ± 17.3 | <0.001 |
| Sex |  |  |  | 0.017 |
| Female | 29 (41%) | 19 (56%) | 10 (28%) |  |
| Male | 41 (59%) | 15 (44%) | 26 (72%) |  |
| BMI (kg/m<sup>2</sup>) | 27.4 ± 4.6 | 26.8 ± 5.0 | 28.3 ± 4.0 | 0.127 |
| Diabetes Mellitus | 21 (30%) | 13 (38%) | 8 (22%) | 0.144 |
| Coronary Artery Disease | 39 (56%) | 17 (50%) | 22 (61%) | 0.350 |
| hs-Troponin T (ng/L) | 35.2 ± 35.2 | 33.5 ± 34.3 | 92.1 ± NA | 0.235 |
| NT-proBNP (pg/mL) | 1,623.8 ± 1,751.4 | 1,805.6 ± 1,939.2 | 1,169.5 ± 1,097.3 | 0.356 |
| Aortic Valve Area (cm<sup>2</sup>) | 0.9 ± 0.2 | 0.8 ± 0.2 | 1.5 ± 0.0 | 0.021 |
| Mean Pressure Gradient (mmHg) | 37.8 ± 13.6 | 39.3 ± 13.0 | 23.2 ± 11.9 | 0.061 |
| ACE-I / ARB Therapy | 25 (36%) | 15 (44%) | 10 (28%) | 0.154 |

<sup>1</sup> Mean ± SD; n (%) <sup>2</sup> Mann-Whitney U; Fisher's exact test

### Widespread immunologic reprogramming occurs via CpG methylation in AVS (Figure 2A-D)

Unsupervised clustering of the 5,000 most variable CpG loci revealed a dominant epigenomic axis that segregated AVS from control circulating T cells along the first dimension, accounting for 15.8% of total variance (*P* = 3.9 × 10^-6^; **Figure 2A**). Genomic context analysis demonstrated that these variable and disease-associated CpGs were not randomly distributed across the genome. Rather, differentially methylated loci were disproportionately enriched within CpG islands and their flanking shores compared with the array background, indicating preferential remodeling of promoter-proximal and regulatory CpG-dense regions (**Figure 2B-C**).

**Figure 2.**
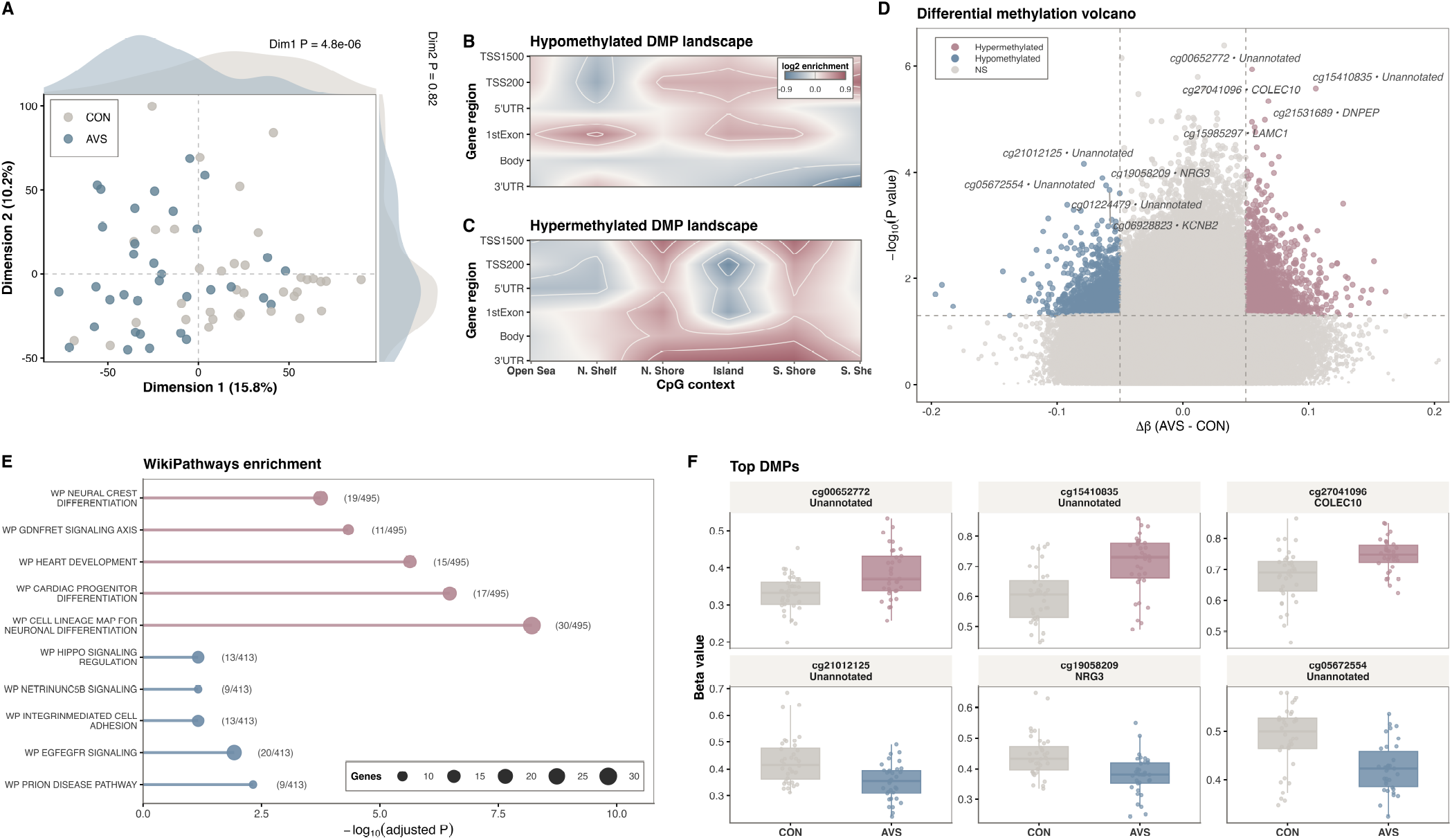
CpG methylation analysis identifies a distinct epigenomic signature in AVS. (A) Unsupervised analysis via multidimensional scaling plot of the top 5,000 most variable CpG sites, with single-dimension density plots quantifying group-wise distribution of AVS and CON via Wilcoxon statistics. Contour plots illustrating genomic and CpG enrichment relative to the Infinium EPIC array background distribution for (B) hypo-methylated and (C) hyper-methylated DMPs (nominal P < 0.05, |Δβ| > 5%). (D) Volcano plot of promoter-associated DMPs highlighting the top five DMPs by P-value. (E) Gene set enrichment of promoter-associated DMPs using the WikiPathways 2022 database, showing hyper-methylated (red) and hypo-methylated (cyan) gene sets. (F) Bar graphs showing the top five most hyper-methylated and hypo-methylated DMPs by effect size.

After adjustment for CAD, age and sex via linear modeling, we identified 3,950 differentially methylated positions (DMPs) at *P* < 0.05 and |Δβ| > 5%, including 1,889 hyper-methylated and 2,061 hypo-methylated loci in AVS (**Figure 2D**). Functional pathway enrichment of DMP-associated genes via gene ontology-term analysis overrepresented aortic valve morphogenesis (15/38, *P* = 6.0 x 10^-10^) and cell-cell adhesion pathways (22/171, *P* = 9.5 x 10^-5^), consistent with coordinated immunologic remodeling at the epigenomic level (**Figure 2E**). CpG-specific candidate searching identified the most statistically significant CpG sites mapped to genes implicated in T cell activation, inflammatory signaling, and immune synapse formation, highlighting discrete regulatory nodes with large effect sizes that may contribute to altered immune cell phenotype in AVS (**Figure 2F**). Taken together, these data support the hypothesis that AVS is accompanied by promoter-enriched epigenomic remodeling of circulating T cells that converges onto immune activation and pro-inflammatory signals.

### Multi-omics factor analysis identifies functional gene regulatory programs in AVS (Figure 3)

To determine whether genome-wide CpG methylation encodes the coordinated, functionally relevant reprogramming of gene expression, we integrated CpG methylation analysis with paired RNA-sequencing data from 54 patients (25 CON, 29 AVS) using multi-omics factor analysis (MOFA). This unsupervised approach identified latent factors capturing shared variance between the transcriptomic and methylomic dimensions. We found that Factor 2 best reflected AVS vs. CON (**Figure 3A**, *P* = 0.0018). Factor 2 strongly correlated with disease status even after adjustment for age and sex (*P* = 0.00024), further supporting its independent representation of AVS disease status (**Figure 3B**).

**Figure 3.**
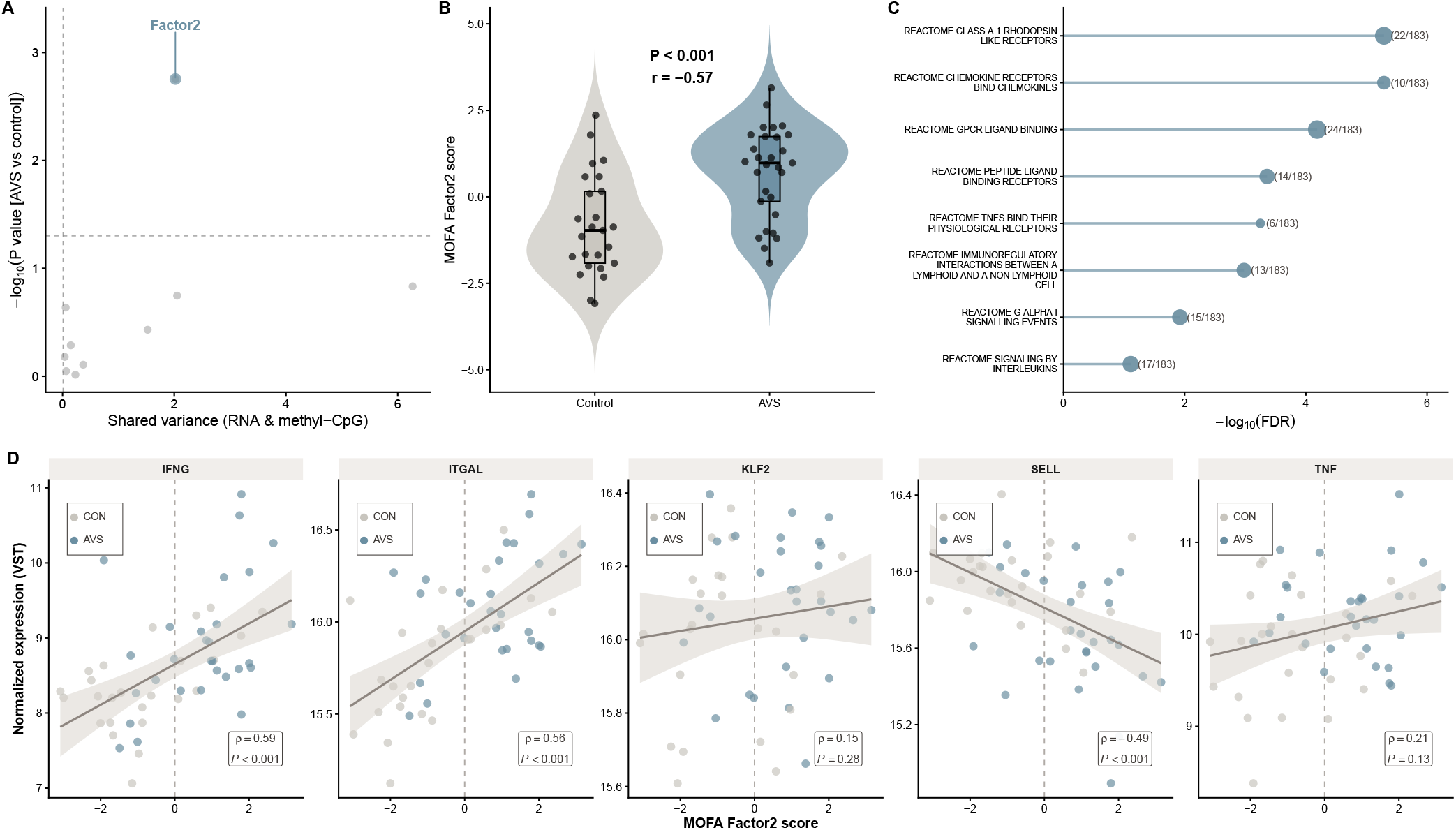
Multi-omics factor analysis (MOFA) identifies a cross-omic, disease-associated latent factor linked to shear-responsive inflammatory programs. (A) Unsupervised MOFA integrating RNA-seq (variance-stabilized counts) and CpG methylation (M-values) across mutual samples. Each point represents a latent factor, plotted by minimum shared variance explained across RNA and methylation (x-axis) versus -log10(BH-adjusted P-value) for association with AVS versus control (y-axis). Factor 2 met prespecified criteria for cross-omic contribution and significant disease association. (B) Distribution of Factor 2 scores in control and AVS samples using violin plot (Wilcoxon P < 0.001). (C) Functional enrichment of genes with the highest positive and negative RNA loadings for Factor 2 using the Reactome 2022 database; bubble size denotes combined enrichment score. (D) Scatterplots of normalized gene expression as a function of Factor 2 score for shear stress and inflammatory mediators, with Spearman correlation coefficients and P-values.

Gene set enrichment analysis of the constituent genes for Factor 2 identified disproportionate immunologic and receptormediated signaling pathways, including chemokine receptor signaling (10/56 enriched, *P* = 1.1 x 10^-6^), TNF ligand-receptor interactions (6/29 enriched, *P* = 6.90 x 10^-5^), and immunoregulatory interactions between lymphoid and non-lymphoid cells (11/123 enriched, *P* = 2.65 x 10^-4^) (**Figure 3C**). At the targeted single-locus level, canonical shear stress-responsive inflammatory mediators integrin subunit alpha L (*ITGAL*), selectin L (*SELL*), tumor necrosis factor (*TNF*), and interferon gamma (*IFNG*) showed robust correlations between expression and the Factor 2 score; by contrast, Krüppel-like factors 1 and 2 (*KLF1, KLF2*) exhibited no association, supporting prior reports that T cells exhibit divergent shear stress-dependent transcriptional reprogramming relative to endothelial cells (**Figure 3D**).^10,22^ Taken together, these observations continue to highlight a functional immunoregulatory T cell reprogramming in AVS encoded by CpG methylation.

### CpG methylation as a clinical risk predictor of aortic stenosis severity

Having identified shear stress-linked epigenetic alterations in circulating T cells, we sought to determine whether an immuno-epigenomic signature could be translated into a probabilistic methylation risk score (MRS). Rather than training a binary case-control classifier, which is vulnerable to confounding by age and other exposures, we modeled continuous hemodynamic severity directly within the AVS cohort. Multivariable linear regression for mean transvalvular gradient was constructed against available clinical covariates (age, sex, smoking status, body mass index, beta-blocker use, and ACE-inhibitor/ARB use) to derive a residualized severity phenotype, ensuring that downstream CpG selection captured shear-stress-attributable variance rather than age- or pharmacotherapy-confounded structure. Probe-wise Spearman correlation was used as a univariate filter, and the top 100 CpGs were carried forward into elastic-net Gaussian regression (α = 0.5; λ = λ_1_se by 5-fold cross-validation), yielding 18 non-zero CpG features constituting the final Severity MRS (**Figure 4A**). Selected loci spanned both hyper- and hypo-methylated coefficients and mapped to genes with established roles in T cell activation and chromatin remodeling, including HDAC9, SOX14, and PRDM15.

**Figure 4.**
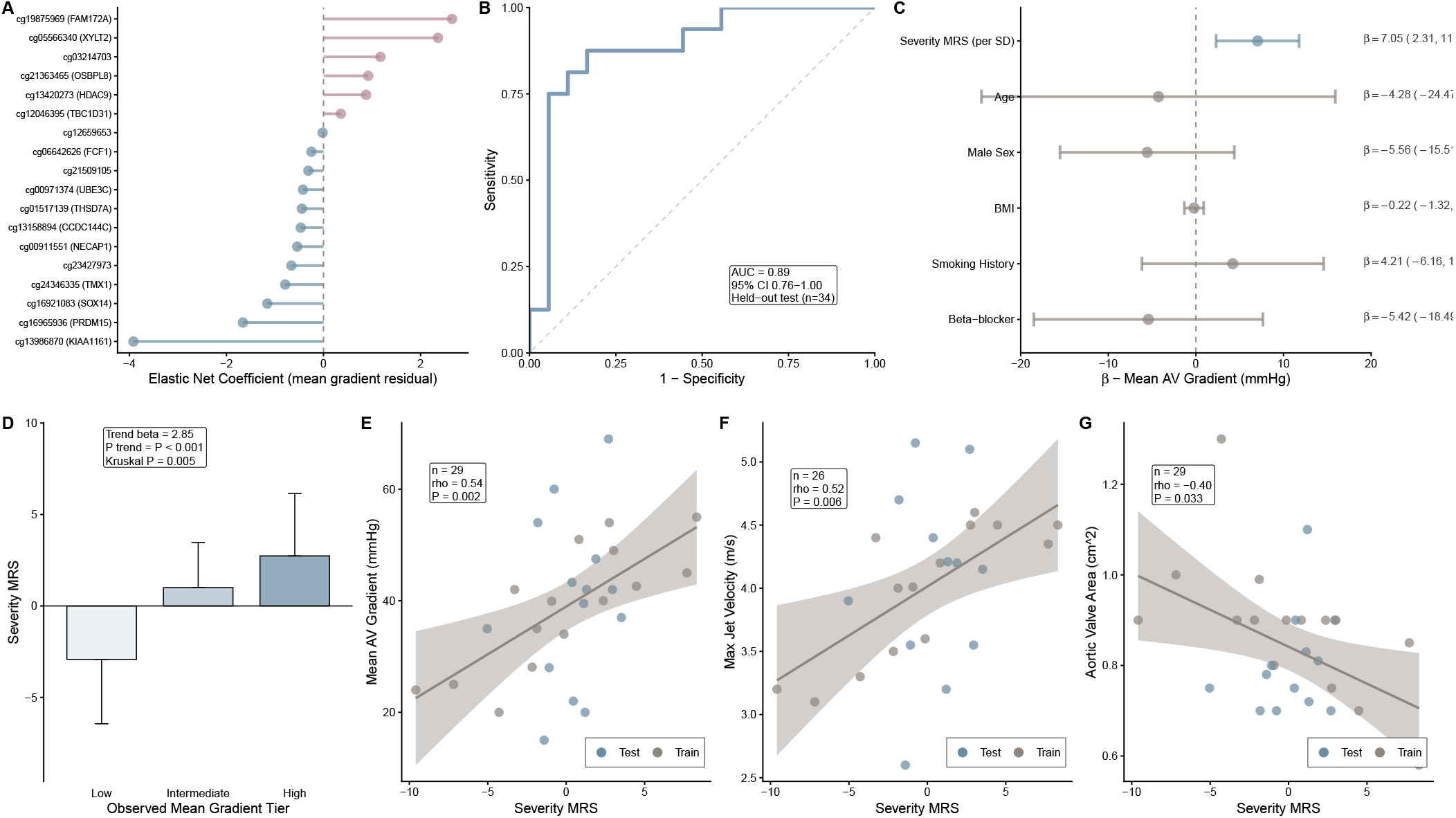
Methylation severity score (MRS) derived from circulating T-cell CpG methylation tracks AVS hemodynamic burden. (A) Lollipop plot of the 18 CpGs retained in the elastic-net Gaussian regression model (α = 0.5; λ = λ1se from 5-fold cross-validation). Points represent non-zero coefficients for mean aortic valve gradient residual; color denotes direction of association. (B) Receiver Operator Curve for the MRS in the projected held-out case-control testing set; AUC = 0.89 (95% CI 0.76-1.00). (C) Multivariable forest plot of mean AV gradient predictors including MRS, age, sex, BMI, smoking history, and beta-blocker therapy; MRS β = 7.05 (95% CI 2.31-11.79). (D) Distribution of MRS by observed mean AV gradient tertile; trend P < 0.001, Kruskal-Wallis P = 0.005. (E) Spearman correlation of MRS with mean AV gradient (ρ = 0.54; P = 0.002; n = 29). (F) Spearman correlation of MRS with maximum jet velocity (ρ = 0.52; P = 0.006; n = 26). (G) Spearman correlation of MRS with aortic valve area (ρ = -0.40; P = 0.033; n = 29).

To evaluate whether a score derived to track severity would also discriminate disease status, we projected the Severity MRS into the full 70-sample case-control dataset and assessed performance in a stratified, held-out test split. The MRS achieved an AUC of 0.89 (95% CI 0.76-1.00; n = 34 held-out subjects) for AVS-versus-control discrimination (**Figure 4B**). Because the model was trained on hemodynamic severity within the AVS cohort and not on diagnostic labels, this estimate should be interpreted as a hypothesis-generating biomarker signal rather than a validated diagnostic claim. External replication in independent, multiethnic cohorts is required before clinical translation.

To define the independent contribution of the MRS beyond clinical covariates, we performed multivariable linear regression of mean transvalvular gradient against the MRS (per SD), age, sex, body mass index, smoking history, and beta-blocker use. The MRS remained the only significant independent predictor of mean gradient (β = 7.05 mmHg per SD; 95% CI 2.31-11.79), outperforming traditional clinical covariates, none of which retained significance after MRS adjustment (**Figure 4C**). Stratification by observed gradient tertile demonstrated a monotonic dose-response relationship, with Severity MRS values increasing progressively from low-to high-gradient groups (trend P < 0.001; Kruskal-Wallis *P* = 0.005; **Figure 4D**).

Across orthogonal echocardiographic indices of AVS severity, the MRS exhibited consistent and significant correlations: with mean transvalvular gradient (Spearman ρ = 0.54; *P* = 0.002; n = 29; **Figure 4E**), maximum aortic jet velocity (ρ = 0.52; *P* = 0.006; n = 26; **Figure 4F**), and inversely with aortic valve area (ρ = −0.40; *P* = 0.033; n = 29; **Figure 4G**). The convergence of significant associations across three independent hemodynamic readouts argues against a chance finding and supports the MRS as a quantitative epigenomic readout of cumulative shear stress exposure on circulating T cells.

### NFAT signaling is a transcriptional nexus of pathologic shear stress in AVS

Having established that the AVS-associated methylome encodes a shear-responsive, immunoregulatory program in circulating T cells, we next sought a candidate signaling axis through which mechanical forces could be transduced into the observed transcriptional state. NFAT is established as a Ca^2+^/TCR-coupled regulator of T cell fate that promotes activation through context-dependent transcriptional partners,^23^ and prior work has documented both NFAT-dependent signaling at low shear stress (0.5-5 dyn/cm^2^) and robust NFAT activation downstream of CD3/CD28-mediated TCR engagement.^10^ Therefore, to test whether HSS is sufficient to induce NFAT nuclear translocation and target gene activation, we subjected Jurkat cells to 20 dyn/cm^2^ using an *in vitro* microfluidics system (ibidi) as a fourfold increase above the physiological transvalvular range (1-5 dyn/cm^2^).^24^ Three hours of 20 dyn/cm^2^ was sufficient to augment NFAT protein nuclear translocation among CD3/CD28 activated T cells (*P* = 2.9 x 10^-2^) to levels comparable to the trans-valvular shear stresses observed in human AVS (**Figure 5B**).^25^ Consistent with its nuclear accumulation, CD3/CD28-stimulated cells exposed to HSS exhibited significant transcriptional induction of canonical NFAT target genes, including IL-2 (*P* = 6.8 x 10^-3^) and IFN-γ (P = 1.2 x 10^-2^); TNF-α (*P* = 0.77) and CCND1 (*P* = 0.08) were not significantly upregulated relative to static conditions (**Figure 5C-F**). Together, these data implicate NFAT as both a TCR-gated and HSS-responsive regulator of T-cell activation that links the hemodynamic stress of AVS to the pro-inflammatory T cell phenotype encoded by the AVS methylome.

**Figure 5.**
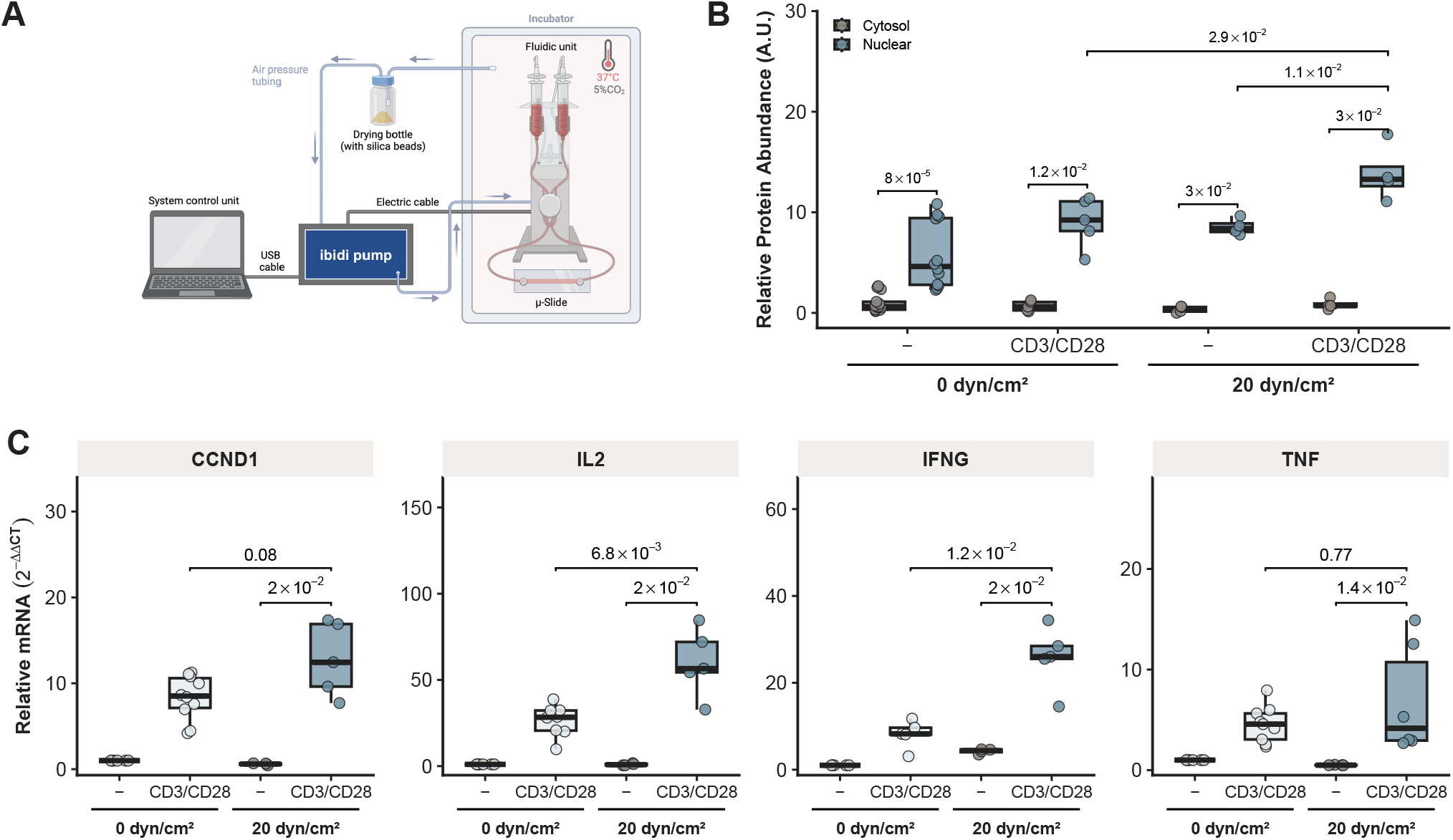
Aortic valvular stenosis-dependent T cell activation of NFAT signaling in vitro. (A) Schematic illustrating the in vitro model for high shear stress (HSS) of Jurkat cells using parallel-flow chamber. Image created using www.biorender.com. (B) NFAT protein expression measured via JessTM with subcellular fractionation between cytosolic (gray) and nuclear (blue) compartments of Jurkat cells in vitro following 1 hour shear stress (0 dyn, 20 dyn) in the presence and absence of CD3 and CD28 antibody stimulation (+CD3/CD28). (C-F) Barplots showing NFAT target gene expression via qPCR in Jurkat cells: (C) CCND1, (D) IL2, (E) IFNG, (F) TNF. Statistical comparisons by Wilcoxon rank-sum test shown as mean ± SD across ≥ 4 independent experiments.

## Discussion

Calcific aortic valve stenosis (AVS) has historically been framed as a degenerative consequence of aging, yet accumulating evidence reframes it as an actively immunomodulated disease in which adaptive immune dysregulation propels progression.^9,26,27^ Pathophysiological parallels with atherosclerosis are striking, including a convergence on infiltrating immune cells to perpetuate tissue damage.^28,29^ We extend this view by identifying a coherent CpG methylation signature within the circulating T cell fraction of patients with AVS, which encodes the transcriptional response to pathological shear stress and T cell activation. Through elastic net regression, we further demonstrate the early clinical potential of this signature as a diagnostic biomarker for AVS.

We present the first genome-wide methylome of circulating T cells in AVS and find a coherent, promoter-enriched signature that segregates disease status and encodes a coordinated pro-inflammatory transcriptional program. Multi-omics integration links this signature to shear-stress-dependent signaling that governs T cell adhesion, migration, and cytokine effector function. That such a stable epigenomic mark is detectable in the circulating compartment, rather than only in valve-infiltrating cells, has a non-obvious explanation: in the trained-immunity paradigm, transient stimuli leave durable, heritable epigenetic imprints on innate immune cells through DNA methylation and histone modifications that persist across cell divisions.^30-32^ Although trained immunity has been characterized predominantly in monocytes and macrophages, analogous chromatin priming has been demonstrated at lymphoid loci,^33^ providing a plausible mechanism by which intermittent hemodynamic exposure could leave a stable methylation footprint on peripheral T cells. Our findings extend this framework to AVS and to the adaptive arm of immunity, building on the multi-omics view of calcific aortic valve disease,^11,23^ and on a conceptual model characterizing adaptive immunity as an active driver of valvular calcification.^24^ We provide direct evidence that circulating T cells, not merely tissue-infiltrating cells, are functionally reprogrammed in AVS through shear-stress-dependent mechanisms, situating the peripheral compartment as both biomarker and contributor.

The gene-regulatory programs encoded by this epigenomic remodeling point to a coherent and testable mechanism. MOFA Factor 2 enrichment for chemokine receptor signaling, TNF ligand-receptor interactions, and lymphoid-non-lymphoid immunoregulatory pathways is consistent with an activated, migratory effector T cell phenotype. Piezo1, a mechanosensitive cation channel constitutively expressed on T cells, converts extracellular shear forces into intracellular calcium signals,^10,12,34^ triggering calcineurin-mediated NFAT nuclear translocation and the downstream effector program of cytokine production and clonal expansion. Recent work refines this circuit, identifying PIEZO1 as the dominant channel for stiffness-coupled MTOC reorientation while ORAI1-mediated store-operated calcium entry drives sustained NFAT translocation; the two channels likely act in concert to convert continuous hemodynamic input into a calibrated transcriptional response.^33-36^ Under conditions of chronic stimulation, as would be expected in the persistent hemodynamic environment of severe AVS, partnerless NFAT binding (in the absence of cooperating AP-1) drives a phenotypic shift toward T cell exhaustion, characterized by sustained but progressively dysregulated effector output.^34-37^ Consistent with prior reports, we observed no significant methylation changes at Krüppel-like factor (KLF) loci, the canonical effectors of the endothelial shear stress response.^35-38^ The cellular response to shear is sharply context-dependent: in endothelial cells, laminar shear induces KLF2 and KLF4, suppressing NF-κB-mediated inflammation and preserving vascular homeostasis; in T cells, by contrast, the same hemodynamic input is transduced through Piezo1-calcineurin-NFAT signaling into pro-inflammatory transcriptional programs that may contribute to AVS pathophysiology. This inversion of the canonical protective response may help explain why peripheral immune-cell biology is mechanistically informative for valve disease.

Comparing circulating T cells from AVS patients with those from healthy controls, we found elevated expression of the NFAT-regulated Th1 cytokines IFN-γ, TNF-α, and IL-2, alongside upregulation of Cyclin D1, consistent with S-phase entry and proliferative reprogramming. These downstream effectors map onto established disease biology. In atherosclerosis, IFN-γ promotes plaque initiation through immune-cell recruitment.^39^ IFN-γ and TNF-α activate antigen-presenting cells via MHC II upregulation, perpetuating localized inflammatory cascades.^40^ Within the aortic valve, IFN-γ enhances the antigen-presenting capacity of resident macrophages and promotes monocyte infiltration. TNF-α drives the osteogenic transition of valvular interstitial cells, increasing alkaline phosphatase activity and bone-matrix protein deposition characteristic of calcific remodeling; IL-2 sustains effector T cell activity within the inflamed leaflet.^41,42^ Together, these data implicate NFAT signaling as a convergence point through which hemodynamic stress in AVS may be transduced into a self-reinforcing pro-inflammatory program in circulating T cells. The therapeutic implication is provocative but must be qualified. Calcineurin inhibitors such as cyclosporine A and tacrolimus block NFAT activation and are clinically established, yet their broad mechanism of action produces nephrotoxicity, hypertension, accelerated atherosclerosis, and increased malignancy risk that argue strongly against their repurposing for AVS. More selective approaches, including the calcineurin-NFAT interaction inhibitor VIVIT, disrupt NFAT activation without compromising calcineurin’s other substrates and have shown promise in preclinical cardiovascular models; whether such targeted modulation could attenuate AVS progression remains untested, and any T cell-directed immunomodulation in an aging population at elevated infection risk demands caution.^43,44^

Beyond mechanism, the methylation signature is clinically actionable. Our parsimonious methylation risk score (MRS) achieved an AUC of 0.89 in the testing cohort, with a monotonic rise in mean transvalvular gradient across MRS tertiles (linear trend P < 0.001, β = 2.85 mmHg per tertile; Kruskal-Wallis P = 0.005), suggesting a continuous, dose response-like relationship between hemodynamic severity and T cell methylation. While these performance metrics require validation in independent cohorts before any clinical claim can be made, they situate the MRS within an emerging body of work demonstrating that blood-based DNA methylation captures cardiovascular risk independently of traditional factors. ARIC investigators showed that leukocyte methylation predicts incident myocardial infarction and coronary heart disease,^45^ and complementary work from the Generation Scotland cohort has shown that DNA methylation-derived epigenetic scores (EpiScores) for circulating proteins augment cardiovascular risk prediction beyond the ASSIGN clinical calculator.^46,47^ Recent epigenome-wide analyses using the CARDIA, Framingham Heart Study, and Generation Scotland cohorts report comparable discriminative performance for cardiovascular outcomes.^47-49^

### Limitations

Several limitations should be acknowledged. First, the cross-sectional design captures a molecular snapshot of established aortic valve stenosis and cannot distinguish whether the observed epigenetic differences precede, accompany, or follow hemodynamically significant disease; their prognostic value therefore remains untested. Second, recruitment from a single German center produced a demographically homogeneous cohort, constraining generalizability of the risk score to multiethnic populations and to patients with bicuspid valve morphology, who were under-represented. Longitudinal, multiethnic validation, ideally extending from pre-clinical aortic sclerosis through post-replacement sampling, will be required to determine whether the methylation signature antedates significant stenosis and whether it regresses after mechanical unloading. Third, although patients with clinically overt inflammatory comorbidities were excluded, residual confounding by subclinical inflammation, age-related immune senescence, or unmeasured pharmacotherapy cannot be ruled out and should be addressed through protocolized adjudication in future cohorts. Fourth, the chromatin-modifying machinery enforcing the hemodynamic shear stress-NFAT-CpG axis in vivo remains uncharacterized. DNMT3A, TET2, and related enzymes govern lineage-specific methylation in memory T cells and gate pro-inflammatory cytokine production.^50-52^ Notably, genes encoding these enzymes are recurrently mutated in clonal hematopoiesis of indeterminate potential, a condition independently linked to cardiovascular inflammation and adverse outcomes.^53-56^ Whether shear-stress-induced NFAT signaling converges on these enzymes through regulated expression, CHIP-driven somatic variation, or both remains unresolved and will require mechanistic studies integrating single-cell methylome and genotype data.

## Conclusion

Severe aortic valve stenosis is accompanied by promoter-enriched epigenomic remodeling of circulating T cells that encodes a coordinated pro-inflammatory program converging on shear stress-dependent NFAT signaling. A parsimonious methylation risk score independently tracks hemodynamic severity, positioning peripheral T cell DNA methylation as a blood-accessible biomarker of valvular disease burden. These findings identify circulating T cells as shear stress-responsive epigenomic proxies of pathological aortic valve disease and provide a foundation for epigenome-guided risk stratification and future investigation of immunomodulatory therapeutic targets in valvular heart disease.

## Abbreviations

AVA: Aortic valve area
AVS: Calcific aortic valve stenosis
CpG: Cytosine-phosphate-guanine
DMP: differentially-methylated position
HSS: High shear stress
MOFA: Multi-omics factor analysis
MRS: Methylation risk score
NFAT: Nuclear factor of activated T cells
TAVI: Transcatheter aortic valve implantation
VIC: Valvular interstitial cell.

## End Notes

### Contributions

Y.Z. designed and supervised the cohort study and microfluidics experiment, performed the experiment and drafted the manuscript. J.L.K., F.S., S.Z., M.C., C.W. and S.F. helped perform the experiments, read and approved the submitted manuscript. M.E.P. is the guarantor of this work and thus accepts responsibility for its computational integrity.

## Acknowledgments

We thank the LIFE & BRAIN GmbH (Bonn, Germany) for their expert support in epigenetic analysis and RNA-Sequencing.

## References

1. Vahanian A, Beyersdorf F, Praz F, et al. 2021 ESC/EACTS Guidelines for the management of valvular heart disease. Eur Heart J. 2022;43:561–632.

2. Coffey S, Roberts-Thomson R, Brown A, et al. Global epidemiology of valvular heart disease. Nat Rev Cardiol. 2021;18:853–864.

3. Thaden JJ, Nkomo VT, Enriquez-Sarano M. The global burden of aortic stenosis. Prog Cardiovasc Dis. 2014;56:565–571.

4. Yutzey KE, Demer LL, Body SC, et al. Calcific aortic valve disease: a consensus summary from the Alliance of Investigators on Calcific Aortic Valve Disease. Arterioscler Thromb Vasc Biol. 2014;34:2387–2393.

5. Kraler S, Blaser MC, Aikawa E, Camici GG, Lüscher TF. Calcific aortic valve disease: from molecular and cellular mechanisms to medical therapy. Eur Heart J. 2022;43:683–697.

6. Mazzone A, Epistolato MC, De Caterina R, et al. Neoangiogenesis, T-lymphocyte infiltration, and heat shock protein-60 are biological hallmarks of an immunomediated inflammatory process in end-stage calcified aortic valve stenosis. J Am Coll Cardiol. 2004;43:1670–1676.

7. Lee SH, Choi J-H. Involvement of immune cell network in aortic valve stenosis: Communication between valvular interstitial cells and immune cells. Immune Netw. 2016;16:26–32.

8. Hulin A, Hortells L, Gomez-Stallons MV, et al. Maturation of heart valve cell populations during postnatal remodeling. Development. 2019;146.

9. Qin Z, Bäck M. Advances in single-cell transcriptomics: unraveling the pathogenesis of calcific aortic valve disease. Cardiovasc Res. 2026;121:2838–2859.

10. Hope JM, Dombroski JA, Pereles RS, et al. Fluid shear stress enhances T cell activation through Piezo1. BMC Biol. 2022;20:61.

11. Zhao R, Zhang J, Zhang S, et al. T cell polarization and NFAT activation are stiffness dependent and differentially regulated by the channels PIEZO1 and ORAI1. Sci Signal. 2026;19:eadt9566.

12. Martinez GJ, Pereira RM, Äijö T, et al. The transcription factor NFAT promotes exhaustion of activated CD8^+^ T cells. Immunity. 2015;42:265–278.

13. Baksh S, Widlund HR, Frazer-Abel AA, et al. NFATc2-mediated repression of cyclin-dependent kinase 4 expression. Mol Cell. 2002;10:1071–1081.

14. Jones PA. Functions of DNA methylation: islands, start sites, gene bodies and beyond. Nat Rev Genet. 2012;13:484–492.

15. Li J, Li L, Wang Y, et al. Insights into the role of DNA methylation in immune cell development and autoimmune disease. Front Cell Dev Biol. 2021;9:757318.

16. Krolevets M, Cate VT, Prochaska JH, et al. DNA methylation and cardiovascular disease in humans: a systematic review and database of known CpG methylation sites. Clin Epigenetics. 2023;15:56.

17. Chen J-Q, Heldman MR, Herrmann MA, et al. Absolute quantitation of endogenous proteins with precision and accuracy using a capillary Western system. Anal Biochem. 2013;442:97–103.

18. Harris VM. Protein detection by Simple Western™ analysis. Methods Mol Biol. 2015;1312:465–468.

19. Fortin J-P, Labbe A, Lemire M, et al. Functional normalization of 450k methylation array data improves replication in large cancer studies. Genome Biol. 2014;15:503.

20. Smyth GK. Linear models and empirical bayes methods for assessing differential expression in microarray experiments. Stat Appl Genet Mol Biol. 2004;3:Article3.

21. Pepin ME, Ha C-M, Crossman DK, et al. Genome-wide DNA methylation encodes cardiac transcriptional reprogramming in human ischemic heart failure. Lab Invest. 2019;99:371–386.

22. Dekker RJ, van Soest S, Fontijn RD, et al. Prolonged fluid shear stress induces a distinct set of endothelial cell genes, most specifically lung Krüppel-like factor (KLF2). Blood. 2002;100:1689–1698.

23. Hogan PG. Calcium-NFAT transcriptional signalling in T cell activation and T cell exhaustion. Cell Calcium. 2017;63:66–69.

24. Cheng CP, Herfkens RJ, Taylor CA. Comparison of abdominal aortic hemodynamics between men and women at rest and during lower limb exercise. J Vasc Surg. 2003;37:118–123.

25. Jhun C-S, Newswanger R, Cysyk JP, et al. Dynamics of blood flows in aortic stenosis: Mild, moderate, and severe. ASAIO J. 2021;67:666–674.

26. Raddatz MA, Madhur MS, Merryman WD. Adaptive immune cells in calcific aortic valve disease. Am J Physiol Heart Circ Physiol. 2019;317:H141–H155.

27. Dutta P, James JF, Kazik H, Lincoln J. Genetic and developmental contributors to aortic stenosis. Circ Res. 2021;128:1330–1343.

28. Cinezan C, Magureanu DC, Hiceag ML, et al. Shared risk factors and molecular mechanisms between aortic stenosis and atherosclerosis: A rationale for therapeutic repositioning. Int J Mol Sci. 2025;26:8163.

29. Cinezan C, Rus CB, Ilias IT. Unraveling the link between aortic stenosis and atherosclerosis: What have we learned? Medicina (Kaunas). 2025;61:1375.

30. Netea MG, Joosten LAB, Latz E, et al. Trained immunity: A program of innate immune memory in health and disease. Science. 2016;352:aaf1098.

31. Saeed S, Quintin J, Kerstens HHD, et al. Epigenetic programming of monocyte-to-macrophage differentiation and trained innate immunity. Science. 2014;345:1251086.

32. Netea MG, Domínguez-Andrés J, Barreiro LB, et al. Defining trained immunity and its role in health and disease. Nat Rev Immunol. 2020;20:375–388.

33. Fanucchi S, Fok ET, Dalla E, et al. Immune genes are primed for robust transcription by proximal long noncoding RNAs located in nuclear compartments. Nat Genet. 2018;51:138–150.

34. Solis AG, Bielecki P, Steach HR, et al. Mechanosensation of cyclical force by PIEZO1 is essential for innate immunity. Nature. 2019;573:69–74.

35. Santana Nunez D, Malik AB, Lee Q, et al. Piezo1 induces endothelial responses to shear stress via soluble adenylyl Cyclase-IP3R2 circuit. iScience. 2023;26:106661.

36. Zheng Q, Zou Y, Teng P, et al. Mechanosensitive channel PIEZO1 senses shear force to induce KLF2/4 expression via CaMKII/MEKK3/ERK5 axis in endothelial cells. Cells. 2022;11:2191.

37. Lan Y, Lu J, Zhang S, et al. Piezo1-mediated mechanotransduction contributes to disturbed flow-induced atherosclerotic endothelial inflammation. J Am Heart Assoc. 2024;13:e035558.

38. Santos F, Sum H, Yan DCL, Brewer AC. Metaboloepigenetics: Role in the regulation of flow-mediated endothelial (dys)function and atherosclerosis. Cells. 2025;14:378.

39. Herrero-Fernandez, Gomez-Bris, Somovilla-Crespo, Gonzalez-Granado. Immunobiology of Atherosclerosis: A Complex Net of Interactions. IJMS. 2019;20:5293.

40. Baratchi S, Zaldivia MTK, Wallert M, et al. Transcatheter Aortic Valve Implantation Represents an Anti-Inflammatory Therapy Via Reduction of Shear Stress-Induced, Piezo-1-Mediated Monocyte Activation. Circulation. 2020;142:1092–1105.

41. Bartoli-Leonard F, Zimmer J, Aikawa E. Innate and adaptive immunity: the understudied driving force of heart valve disease. Cardiovascular Research. 2021:cvab273.

42. Kaden JJ, Kiliç R, Sarikoç A, et al. Tumor necrosis factor alpha promotes an osteoblast-like phenotype in human aortic valve myofibroblasts: a potential regulatory mechanism of valvular calcification. Int J Mol Med. 2005;16:869–872.

43. Aramburu J, Yaffe MB, López-Rodríguez C, Cantley LC, Hogan PG, Rao A. Affinity-driven peptide selection of an NFAT inhibitor more selective than cyclosporin A. Science. 1999;285:2129–2133.

44. Yu H, van Berkel TJC, Biessen EAL. Therapeutic potential of VIVIT, a selective peptide inhibitor of nuclear factor of activated T cells, in cardiovascular disorders. Cardiovasc Drug Rev. 2007;25:175–187.

45. Agha G, Mendelson MM, Ward-Caviness CK, et al. Blood leukocyte DNA methylation predicts risk of future myocardial infarction and coronary heart disease. Circulation. 2019;140:645–657.

46. Gadd DA, Hillary RF, McCartney DL, et al. Epigenetic scores for the circulating proteome as tools for disease prediction. Elife. 2022;11:e71802.

47. Chybowska AD, Gadd DA, Cheng Y, et al. Epigenetic contributions to clinical risk prediction of cardiovascular disease. Circ Genom Precis Med. 2024;17:e004265.

48. Zheng Y, Joyce BT, Hwang S-J, et al. Association of cardiovas-cular health through young adulthood with genome-wide DNA methylation patterns in midlife: The CARDIA study. Circulation. 2022;146:94–109.

49. Westerman K, Fernández-Sanlés A, Patil P, et al. Epigenomic assessment of cardiovascular disease risk and interactions with traditional risk metrics. J Am Heart Assoc. 2020;9:e015299.

50. Perkins B, Novis CL, Baessler A, et al. Dnmt3a-dependent de novo DNA methylation enforces lineage commitment and pre-serves functionality of memory Th1 and Tfh cells. bioRxivorg. 2025.

51. Zhang X, Su J, Jeong M, et al. DNMT3A and TET2 compete and cooperate to repress lineage-specific transcription factors in hematopoietic stem cells. Nat Genet. 2016;48:1014–1023.

52. Zhong C, Zhu J. Tet2: breaking down barriers to T cell cytokine expression. Immunity. 2015;42:593–595.

53. Abplanalp WT, Cremer S, John D, et al. Clonal hematopoiesis-driver DNMT3A mutations alter immune cells in heart failure. Circ Res. 2021;128:216–228.

54. Cobo I, Tanaka T, Glass CK, Yeang C. Clonal hematopoiesis driven by DNMT3A and TET2 mutations: role in monocyte and macrophage biology and atherosclerotic cardiovascular disease. Curr Opin Hematol. 2022;29:1–7.

55. Nadar S, Dohadwala TK, Kumaresan N, Ahamed SI, Fatima S. Clonal hematopoiesis at the crossroads of Inflammaging and car-diovascular disease: Mechanistic insights and translational horizons. Clin Hematol Int. 2025;7:54–63.

56. Mohammed Ismail W, Fernandez JA, Binder M, et al. Single-cell multiomics reveal divergent effects of DNMT3A-and TET2-mutant clonal hematopoiesis in inflammatory response. Blood Adv. 2025;9:402–416.

