## Supplemental Figure S1 for "Aortic valve stenosis promotes pathological shear stress-dependent epigenomic dysregulation in circulating T cells"

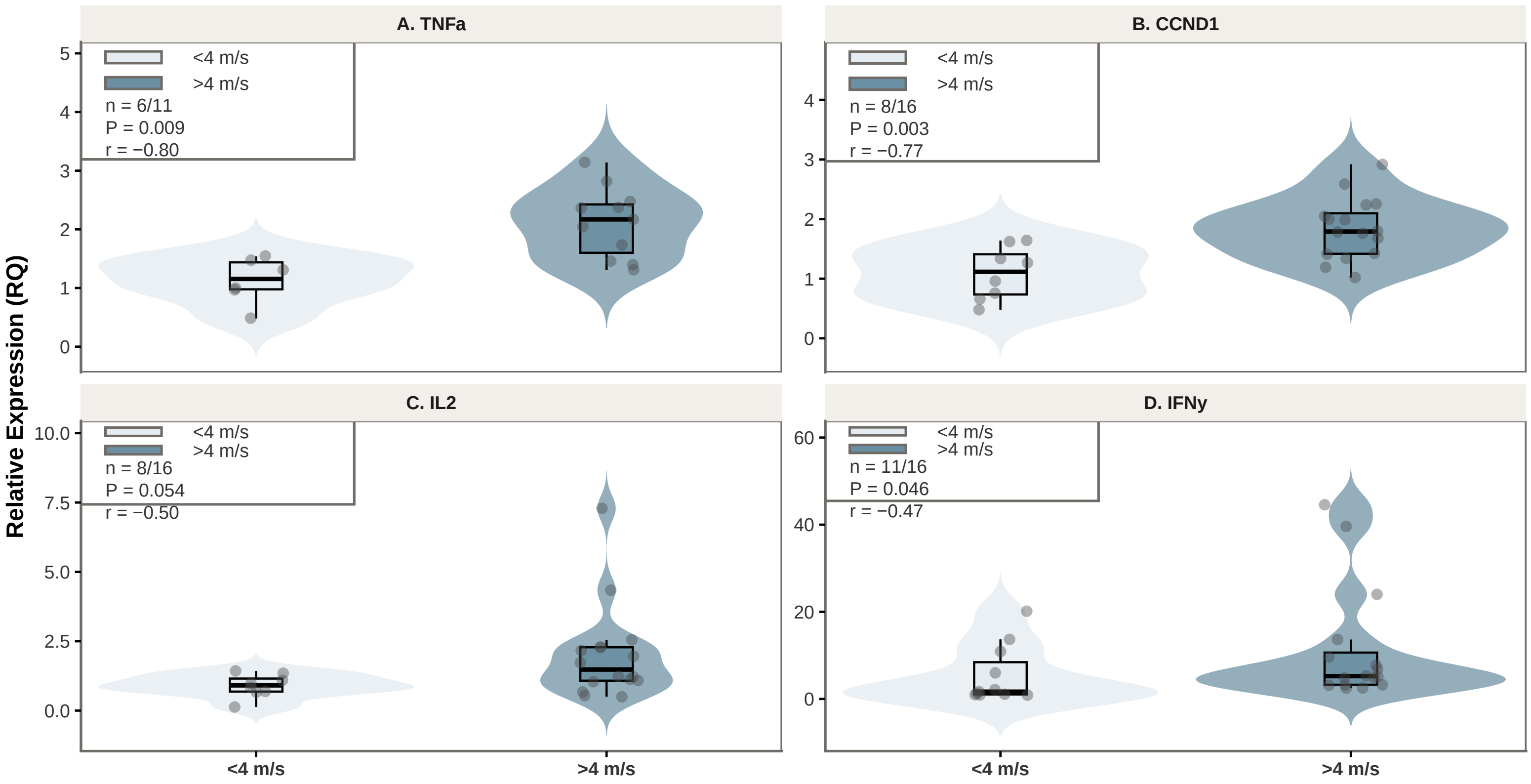


**Supplemental Figure S1.**

Quantitative PCR expression of NFAT−associated genes stratified by peak aortic jet velocity (<4 m/s vs >4 m/s). Violin plots show the distribution of relative expression (RQ), box plots show median and interquartile range, and points represent individual samples. Two−sided P values were calculated using the Wilcoxon rank−sum test; r denotes the rank−biserial effect size.
