## Supplementary figures and images for "Aortic valve stenosis promotes pathological shear stress-dependent epigenomic dysregulation in circulating T cells"

### Supplemental Figure S2

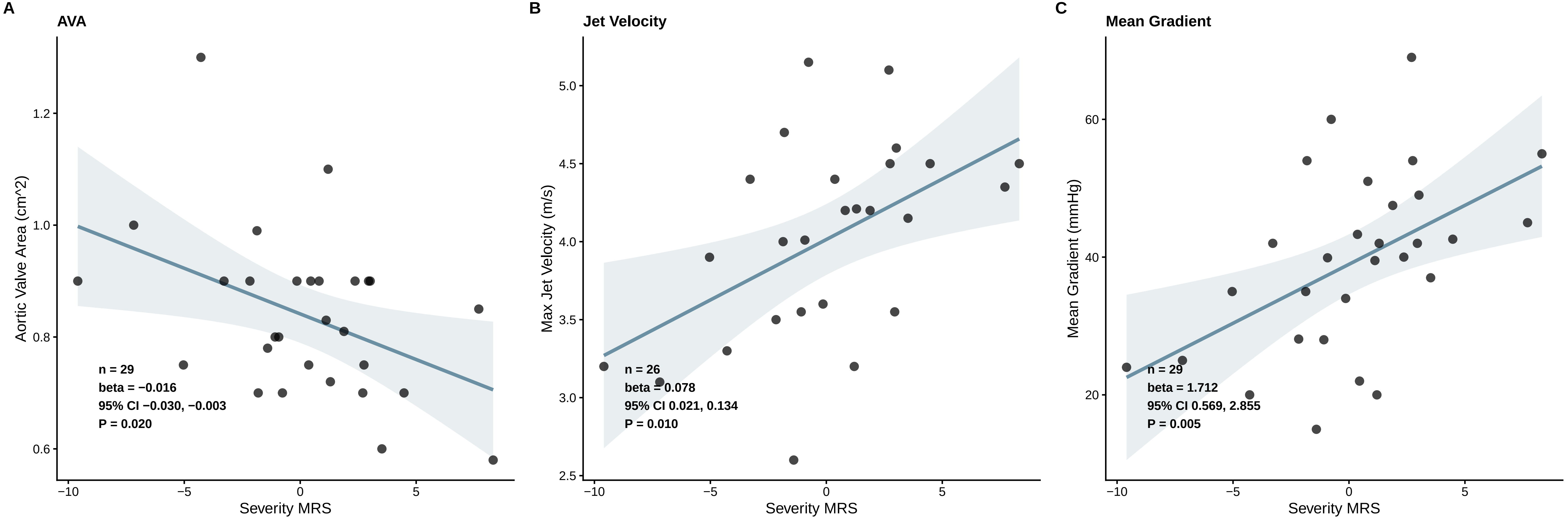

### Supplemental Figure S3

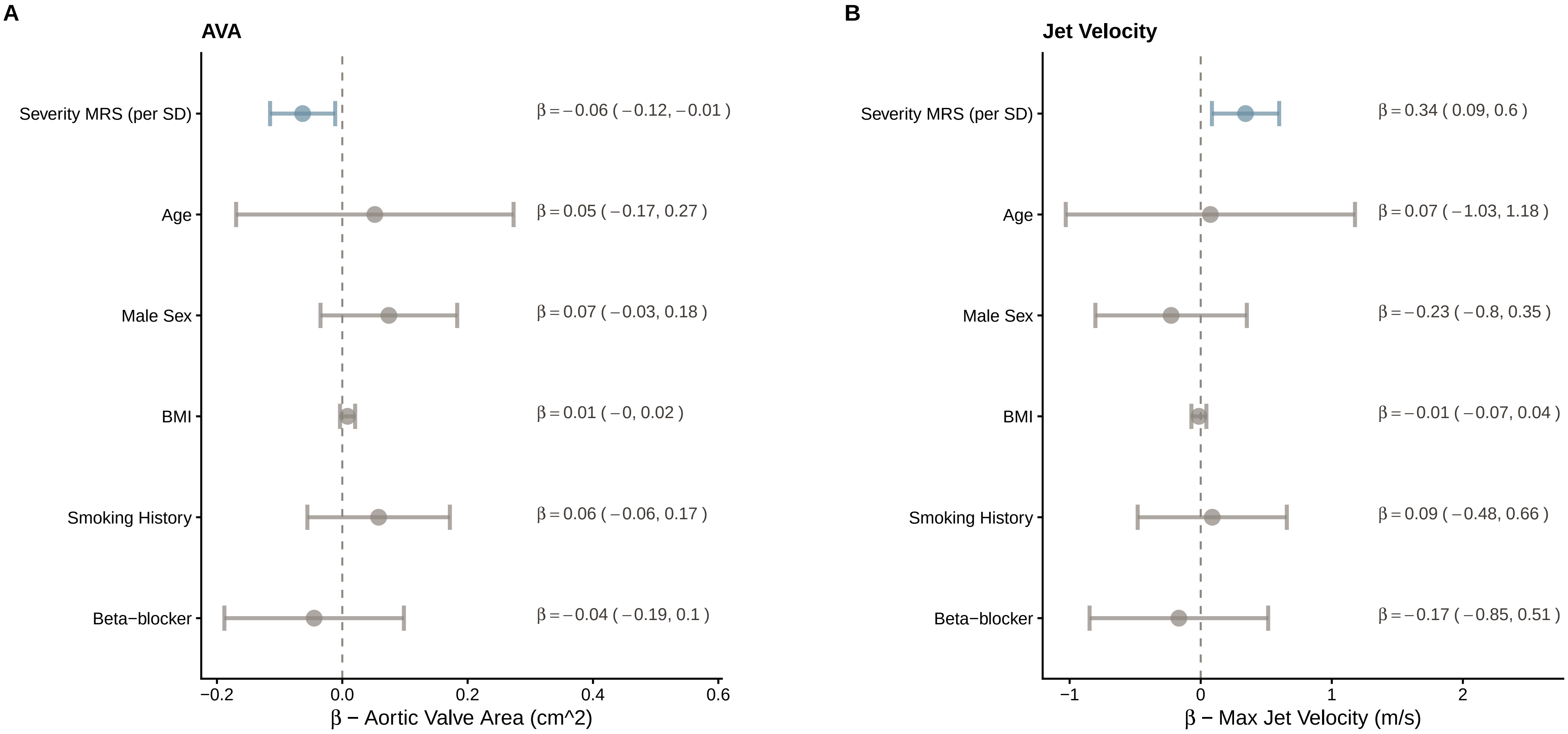

### Supplemental Figure S4

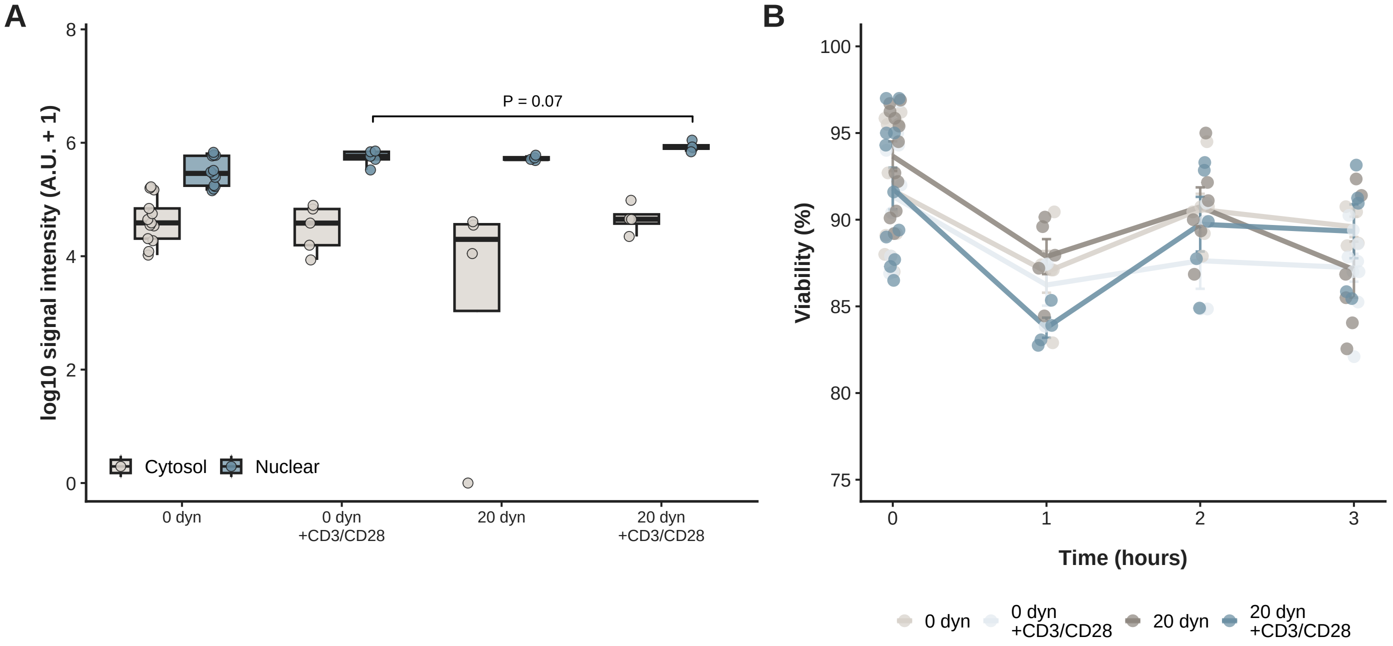

### Supplemental Figure S5

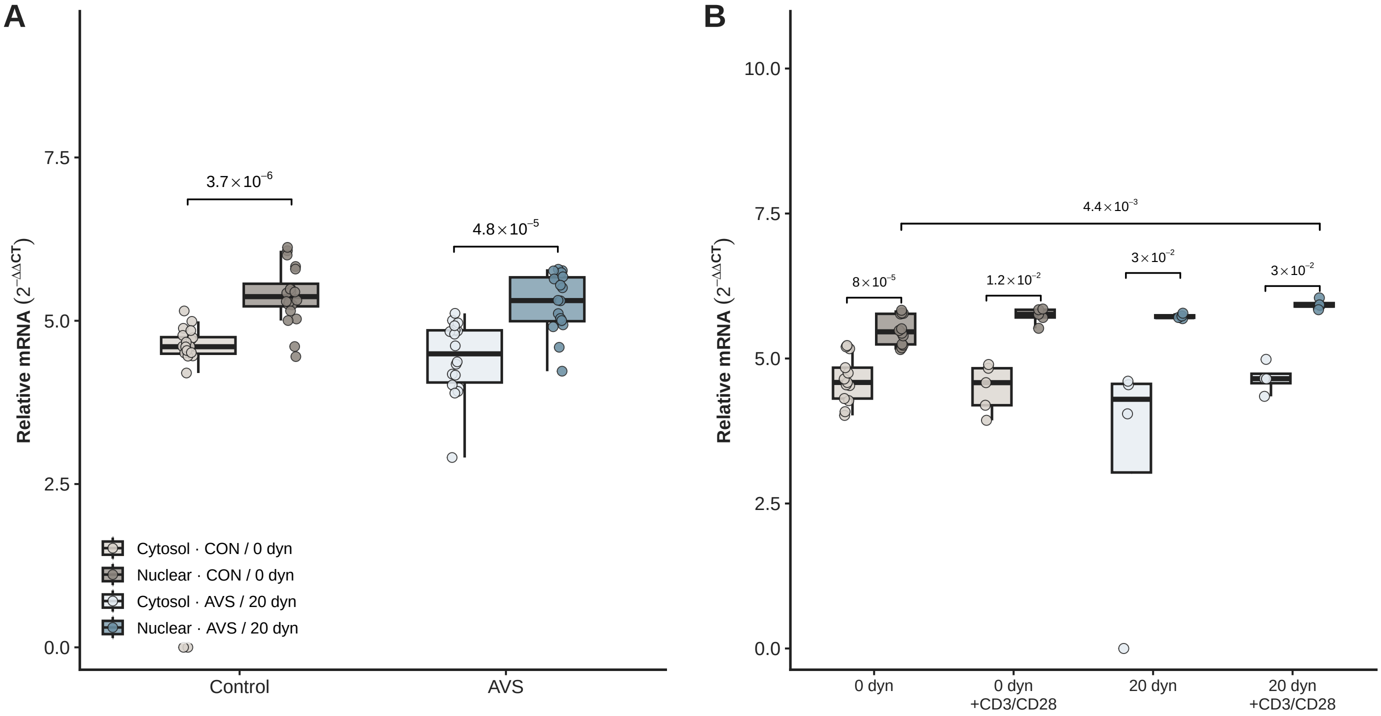
